# Visualizing sarcomere and cellular dynamics in skeletal muscle to improve cell therapies

**DOI:** 10.1101/2024.02.01.578471

**Authors:** Judith Hüttemeister, Franziska Rudolph, Michael H. Radke, Claudia Fink, Dhana Friedrich, Stephan Preibisch, Martin Falcke, Eva Wagner, Stephan E. Lehnart, Michael Gotthardt

## Abstract

The giant striated muscle protein titin integrates into the developing sarcomere to form a stable myofilament system that is extended as myocytes fuse. The logistics underlying myofilament assembly and disassembly have started to emerge with the possibility to follow labeled sarcomere components. Here, we generated the mCherry knock-in at titin’s Z-disk to study skeletal muscle development and remodeling. We find titin’s integration into the sarcomere tightly regulated and its unexpected mobility facilitating a homogenous distribution of titin after cell fusion – an integral part of syncytium formation and maturation of skeletal muscle. In adult mCherry-titin mice, treatment of muscle injury by implantation of titin-eGFP myoblasts reveals how myocytes integrate, fuse and contribute to the continuous myofilament system across cell boundaries. Unlike in immature primary cells, titin proteins are retained at the proximal nucleus and do not diffuse across the whole syncytium with implications for future cell-based therapies of skeletal muscle disease.

## Introduction

During skeletal muscle development, the first myogenic wave starts around E11 with the fusion of embryonic myoblasts at the limb buds and the dermomyotome and is accomplished by a cascade of myogenic transcription factors like myogenic factor 5 (Myf5) and myoblast determination protein (MyoD). In the second myogenic phase (E14.5 – E17.5) these primary fibers fuse with fetal myoblasts to build secondary fibers (Chal and Pourquié, 2017). Thereafter, some myoblasts remain less differentiated to become satellite cells, the stem cell pool in adult muscle (Relaix et al., 2005). They enter quiescence a few weeks after birth and subsequently, hypertrophy is the main driver of muscle growth (Chal and Pourquié, 2017). In the adult, satellite cells can get activated to facilitate muscle regeneration with differentiation to myoblasts and then myocytes, which eventually undergo cell fusion to form new fibers and extend existing ones (Almada and Wagers, 2016).

Titin is abundantly expressed in vertebrate striated muscle (Wang et al., 1979), determines skeletal muscle structure and function (Horowits et al., 1986), and is extensively spliced to produce isoforms with differential mechanical properties (Cazorla et al., 2000; Guo et al., 2012; Li et al., 2012). These vary between heart and skeletal muscle and integrate into the Z-disk and M-band of the sarcomere to form a continuous elastic filament system along the myofiber (Gregorio et al., 1998; Obermann et al., 1997). The process is tightly orchestrated (Rudolph et al., 2019) and the resulting scaffold facilitates proper localization of sarcomeric proteins along the filament (Rudolph et al., 2020). Thus, titin has been proposed to act as a molecular ruler and as a blueprint for sarcomere assembly (Tonino et al., 2017).

With the use of fluorescent titin proteins expressed at physiological levels in knock-in mice, we have obtained insights into the titin lifecycle and sarcomere dynamics in cardiomyocytes (da Silva Lopes et al., 2011; Rudolph et al., 2019). In contrast to the heart, skeletal muscle cells form large syncytia, which contain nuclei of several fused cells. How titin moves along the large syncytium and how titins derived from different nuclei within the syncytium are organized and integrated after cell fusion has so far been prohibitively difficult to assess.

Here, we have extended the portfolio of fluorescent titin mice with the fluorophore mCherry inserted into titin’s Z-disk region to follow titin not only around the sarcomere, but also during cell fusion. The combination of the eGFP knock-in mice as cell donor and mCherry knock-in mice as recipients improves the evaluation of cell-based therapies, as we not only learn where injected cells go, but demonstrate reconstitution of the myofilament in regenerating muscle and the limits of delivering healthy protein in a syncytium in cell-based therapy.

## Results

### The Ttn(Z)-mCherry mouse

To follow titin dynamics during cell fusion of skeletal muscle cells we relayed on our established reporter mice, with fluorophores integrated into the M-Band (da Silva Lopes et al., 2011) or Z-disk (Rudolph et al., 2019) region of titin. The knock-in approach resulted in the physiological expression of fluorescent-tagged titin and did not interfere with sarcomere assembly, titin integration, and striated muscle function. To improve the signal intensity of the red fluorophore and thus enable the analysis of skeletal muscle, we replaced dsRed at the Z-disk exon 27, C-terminal of the Z9 domain with mCherry (Fig. 1a, b). The process involved homologous recombination in ES-cells, blastocyst injection and removal of the NEO cassette with FLP recombinase (Fig. 1a). Homozygous and heterozygous Ttn(Z)-mCherry mice assembled functional sarcomeres with intermediate signal intensity in muscles of heterozygous mice (Fig. S1a, b). As expected from models created earlier, there was no obvious adverse phenotype (Rudolph et al., 2019), no difference in heart to bodyweight ratio (Fig. S1c, d), and proper co-localization of the mCherry-fluorophore with the Z-disk protein α-actinin in homozygous Ttn(Z)-mCherry and double-heterozygous Ttn(Z)-mCherry/Ttn(M)-eGFP mice (Fig. 1c). Live imaging of myotubes with the SpinningDisk microscope (Fig. 1d) confirmed an increased signal intensity of Ttn(Z)-mCherry compared with Ttn(Z)-dsRed mice (Fig. 1e). With the improved red fluorescent label at titin’s Z-disk it is now possible study the dynamics of endogenously expressed titin simultaneously at Z-disk and M-Band and even in immature myocytes during cell fusion.

**Figure 1.**
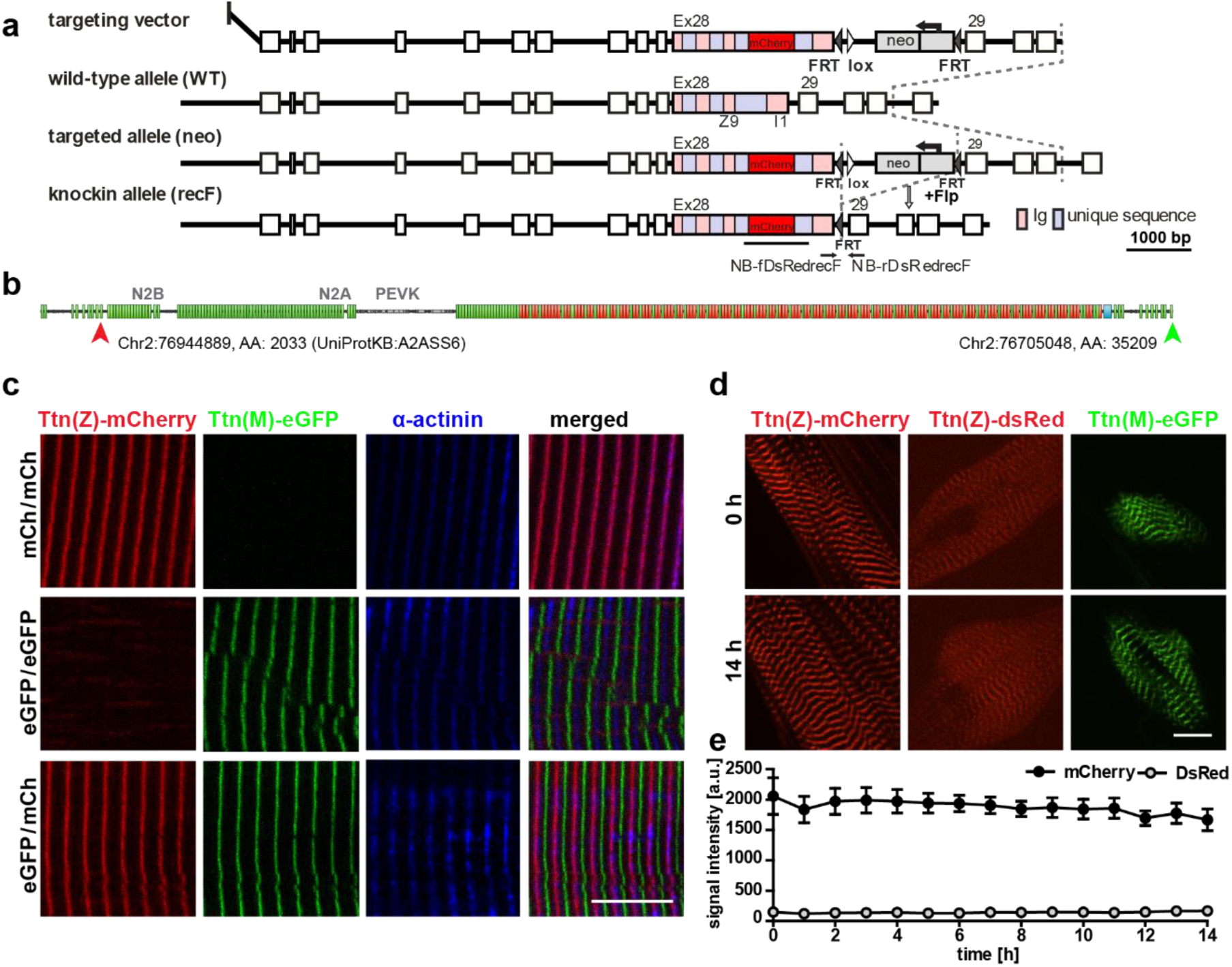
Generation and validation of a titin Z-disk knock-in with enhanced fluorescence. a) Targeting strategy to insert mCherry into titin’s Exon 28 (Z-disk). b) mCherry is integrated outside the Z9 domain (red arrow) and GFP at the c-terminus (green arrow). c) Alternating red and green fluorescent staining in tibialis anterior (TA) of homozygous titin(Z)-mCherry, homozygous titin(M)-eGFP and double-heterozygous mice. Co-staining for α-actinin as a marker of the Z-disk confirms a proper localization of the mCherry fluorophore. d) Simultaneous live imaging of myotubes with dsRed or mCherry fused to titin reveals higher intensity and better signal-to-noise ratio for the mCherry fluorophore. Scale bar 10 µm. e) Stability of the fluorescent signal with minor changes over 14 hours and higher intensity of the mCherry signal.

### Titin kinetics in double-heterozygous myotubes

Measurements of titin kinetics in cardiomyocytes revealed that titin is not a static backbone, but dynamically exchanged in the sarcomere within hours with a faster exchange rate at its Z-disk region (da Silva Lopes et al., 2011; Rudolph et al., 2019). The different cell morphology and titin isoform composition between heart and skeletal muscle prompted the question, whether titin kinetics is different in skeletal muscle cells, which we addressed using fluorescence recovery after photobleaching (FRAP) in Ttn(Z)-mCherry/Ttn(M)-eGFP double-heterozygous myotubes. In the representative images the Ttn(Z)-mCherry signal reemerges already after 1 h as compared to 4 h for the Ttn(M)-eGFP signal and documented in the respective line profiles (Fig. 2a). To confirm that the recovery of the fluorescence signal is due to titin protein exchange and not caused by a reactivation of the fluorophore, we performed the same experiment in fixed cells, where the striated signal pattern did not recover (Fig. S2a). Only minimal background fluorescence was recovered in fixed cells after 14 h with no difference between Ttn(Z)-mCherry and Ttn(M)-eGFP (Fig. S2b). In contrast, there was a significant difference in fluorescence recovery and hence protein exchange between mCherry-labeled Z-disk titin and eGFP-labeled M-Band titin in living cells (Fig. 2b). The mobile fraction of Z-disk titin is significantly higher than the mobile fraction of M-Band titin with 73 vs. 46% (Fig. 2c) - although there is variability between individual cells. The faster recovery of the mCherry-titin signal is also reflected in its significantly reduced exchange half-life of 1.5 h compared to the 4.9 h for the Ttn(M)-eGFP signal (Fig. 2d). To the average fluorescence recovery (Fig. 2b) as well as for the recovery in most individual cells, a two-phase association curve provided a better fit to the data points than the classical one-phase association curve, suggesting that the measured signal can be attributed to two protein isoform populations with different kinetics. The percentage of the fast population is significantly higher for Ttn(Z)-mCherry than for Ttn(M)-eGFP with 37 vs. 16% (Fig. 2e). Quantification of the fluorescence signal at the opposite ends of the half-sarcomere (red signal at the M-Band and green signal at the Z-disk) allowed us to quantify the kinetics of non-integrated titin. Outside their respective integration sites, there was no significant difference anymore between the recovery of mCherry-labelled Z-disk region and the eGFP-labelled M-Band region of titin (Fig. 2f). However, while there was no difference in mobile fraction and ratio of slow to fast population, there was still a significant difference in exchange half-life (Fig. S2f-h). There was no significant difference between integrated and non-integrated Z-disk titin (determined at its integration site and between, respectively), but there was an increased fluorescence recovery of non-integrated titin-eGFP (significant from 6 to 10 h). It appears that titin exchange kinetics in skeletal muscle myotubes are faster at titin’s Z-disk versus its M-Band with similar rates as in embryonic cardiomyocytes (Rudolph et al., 2019), although the cells are structural different and contain different titin isoforms.

**Figure 2.**
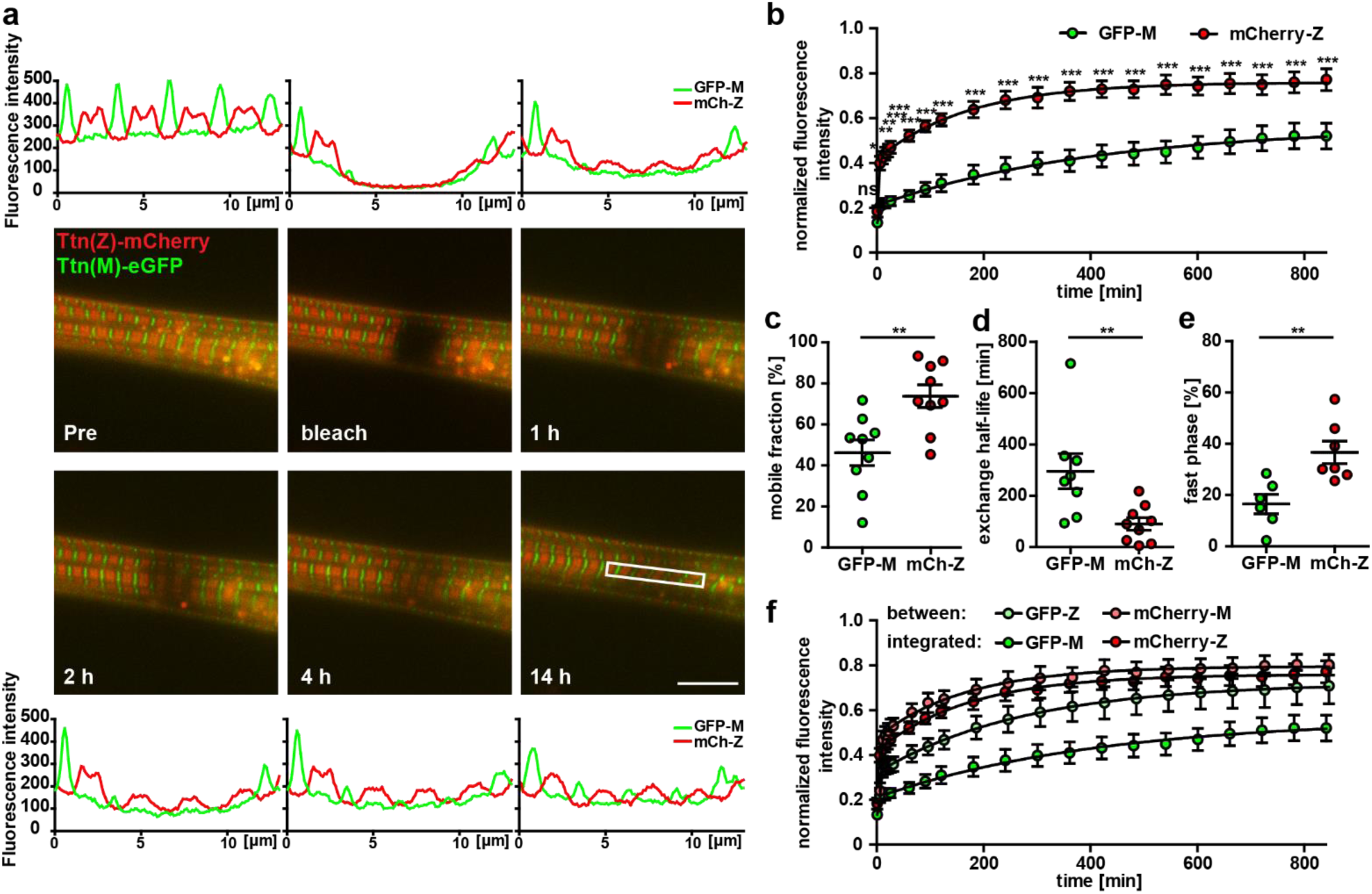
Titin mobility in myotubes at Z-disk and M-band a) Recovery of the sarcomeric titin signal within 14 h. Intensity profiles of the bleached regions (white rectangle). Scale bar 10 µm. b) The mCherry labelled Z-disk titin (mCh-Z) recovers significantly faster than the GFP-labeled M-Band titin (GFP-M; n = 3; 3 cells per experiment, two-way ANOVA) with mobile fraction increased (c) and exchange half-life reduced (d). e) The recovery of fluorescent titin is biphasic with a higher contribution of the fast phase for Z-disk vs. M-band titin. c-e) n = 6 to 9 cells per group, one-way ANOVA for c, d and e. f) Non-integrated GFP-labelled titin (GFP signal outside the M-band) recovers faster than M-band integrated GFP-titin. Samples with obvious decrease in cell quality during imaging where excluded from the analysis.

### Sarcomeric protein dynamics after cell fusion

A remarkable feature of skeletal muscle cells is that they form large, multi-nucleated syncytia arising from cell fusion. It is not completely understood so far, how sarcomeric proteins of different ancestor cells are distributed and integrated along the myotube.

To address these questions, we co-cultured myoblasts of homozygous Ttn(M)-eGFP and homozygous Ttn(Z)-mCherry mice at high density and differentiated them by withdrawing growth factors one day later for 2-3 days to induce their fusion. After fixation, we found cells at different states of differentiation (Fig. 3). In the first phase of fusion, cells had made initial contact as determined by visualizing cell contact formation with M-Cadherin staining (Fig. S3a), but titin-eGFP and mCherry-titin proteins had not mixed yet (Fig. 3a), suggesting that membrane breakdown had not happened. Other cells had already fused as differentially labelled titin had started to mix (Fig. 3b). Here, the alternating mCherry and eGFP signals in the central region of the syncytium indicate the proper integration of titin protein originating from different nuclei. The lower region contained mainly mCherry-titin, suggesting that the lower nucleus originated from a Ttn(Z)-mCherry homozygous myocyte. In some syncytia titin had already distributed completely (likely an early fusion event), so that the nuclei could have originated from either background (Fig. 3c). Sarcomeric proteins such as α-actinin are present (Fig. S3b,c) and localize towards their position in the newly formed sarcomeres throughout the cell (Fig. S3).

**Figure 3.**
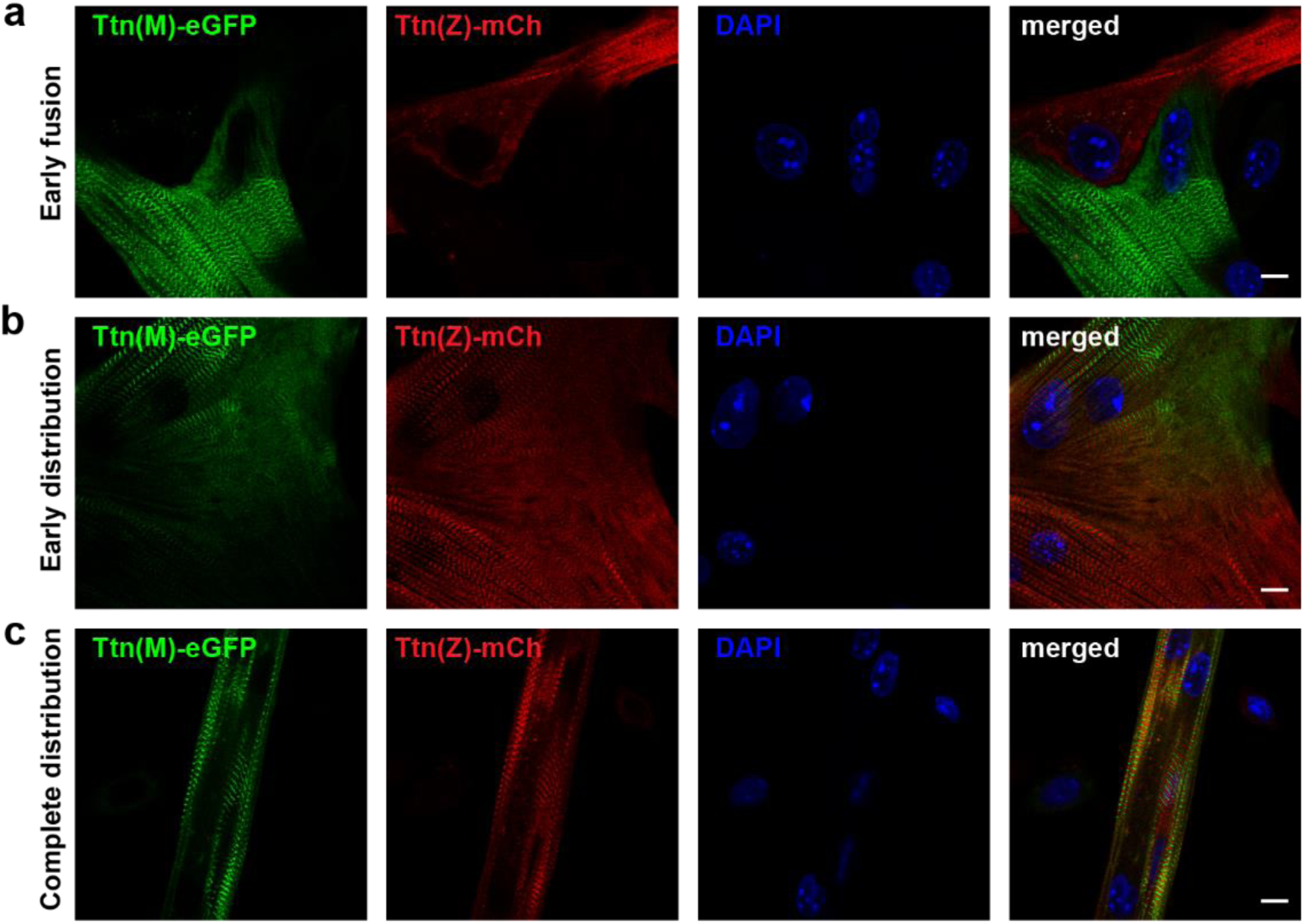
After cell fusion, titin is distributed throughout the myotube. Satellite cells were isolated from homozygous titin(Z)-mCherry and titin(M)-eGFP mice. After co-cultivation and differentiation to myotubes, cells were fixed at different stages of cell fusion, from first contact and early fusion (a) to early (b) and late (c) distribution of titin proteins. Scale bar 10 µm.

### Following titin along the syncytium in real-time

To follow the progression of cell fusion and titin distribution we acquired time lapses from 4-6h after initiating differentiation for 16h total. We successfully recorded several fusion events with myotubes of different sizes fused in different orientations (cell-to-cell or perpendicular). As determined by the sarcomere structure, we documented fusion events between two immature cells or between an immature cell and a mature cell / myotube. We followed the localization of nuclei expressing red or green titin in the syncytium and quantified the distribution of titin over time (Fig. 4). The area where both titin signals were present above threshold levels were subdivided into areas with mainly red signal, mainly green signal and an area with similar amounts of red and green titin. We also provide a movie to follow the fusion event in a timelapse (supplementary Movie S1).

**Figure 4.**
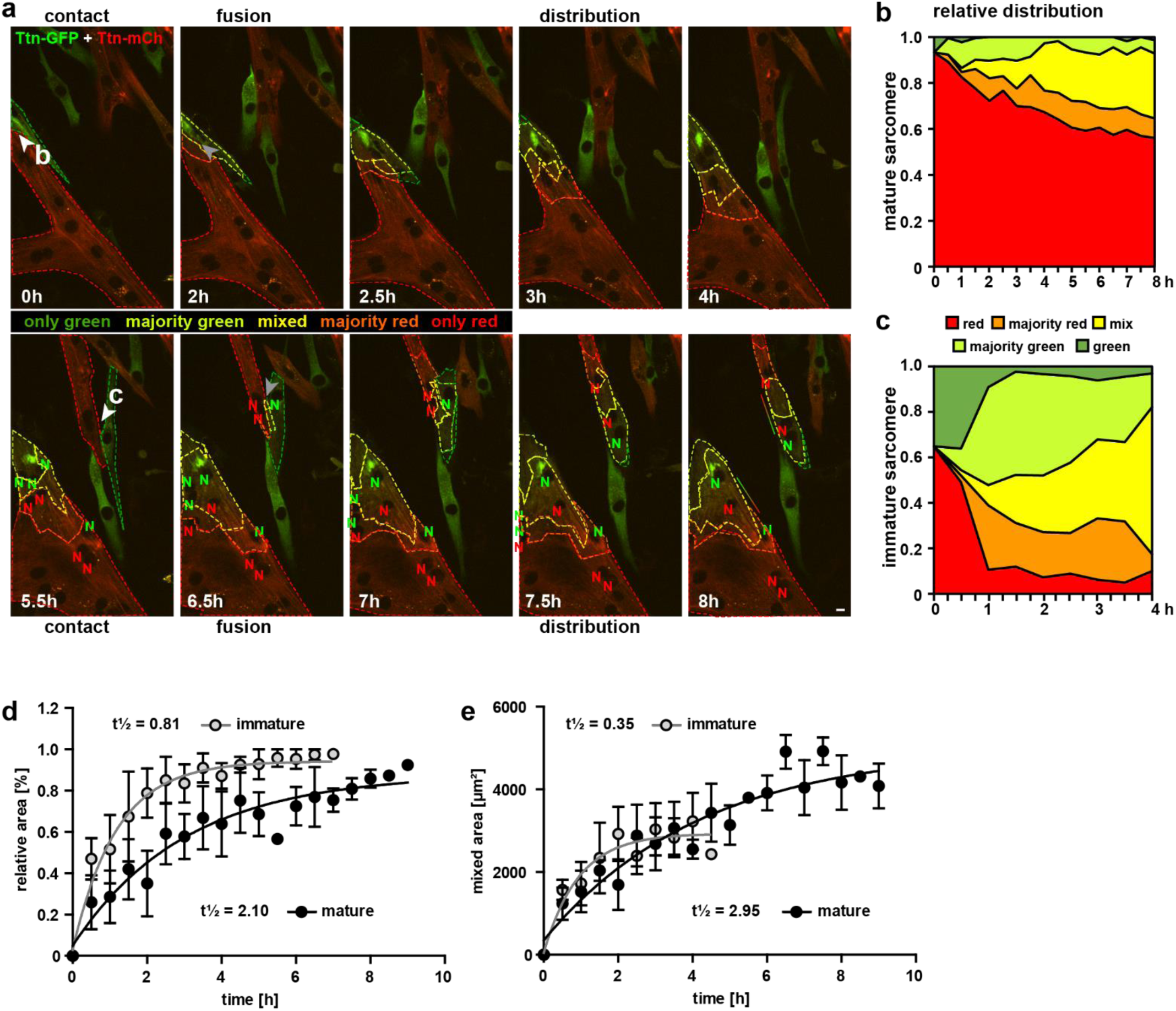
Cell fusion and redistribution of red and green titin in skeletal muscle cells. a) Live imaging of cell fusion of homozygous Ttn(Z)-mCherry and Ttn(M)-eGFP myogenic cells (2 frames per h). Arrows indicate the initiation of cell fusion between a small Ttn-eGFP myocyte and a mature myofiber (b) compared to the fusion of two small, immature cells (c). Nuclei expressing red and green fluorescent titin are labeled with red and green “N”, respectively. Regions with different fluorescent titin ratios are indicated with dashed outlines. Gradient bar (8h) indicates the range of titin spread between two neighboring nuclei. Scale bar 10 µm. b,c) Titin distribution is measured as the area containing red only, majority red, even mix, majority green, or green only based on thresholds set at 20% and 50% maximal fluorescence intensity as detailed in material and methods. b) fusion between a green immature and large red mature cell leads to a gradual redistribution of green titin to less than half of the resulting syncytium within 8 hours. c) Fusion between a red and green immature cell leads to a rapid redistribution within the first hour that is almost compete by 4 hours. Relative (d) and absolute (e) increase of the area with mixed red and green titins in immature cells fusing with mature cells (black) versus immature cells (grey) indicate a >2.5x faster titin distribution when both cells are immature (n = 4 to 5), two-way ANOVA.

In Fig. 4a, two fusion events are indicated with white arrows directed at the points of contact. The first fusion event at 0h of an eGFP myocyte with a large mature multinucleated mCherry myotube leads to the gradual diffusion of eGFP-titin that ultimately contributes to <50% of the sarcomeres (Fig. 4b). The second fusion event at 5.5h, two small immature cells fuse, followed by the rapid distribution of mCherry-titin and titin-eGFP. Within 1h, about 90% of the area is occupied by titins from both original cells (Fig. 4c). For statistical validation of the increased speed of titin distribution in cells fusing to immature versus mature myotubes, we quantified 9 fusion events. To minimize effects of size differences of the syncytia, we excluded very small (< 1000 µm^2^) and very large (> 10,000 µm^2^) cells. As there was still a trend for cells with a mature sarcomere structure to be larger, we provide relative (Fig. 4e) and absolute values (Fig. 4f), with titin mobility (t1/2) reduced by more than two-fold in immature cells undergoing fusion.

### Titin mRNA localization after cell fusion

To dissect the contribution of titin mRNA vs. protein to titin mobility along the syncytium, we visualized titin mRNA originating from different myocytes using smFISH with probes directed against GFP (labelled with Quasar570) and mCherry mRNA (labelled with Quasar670). Homozygous Ttn(Z)-mCherry and Ttn(M)-eGFP cells were plated together and differentiated to induce cell fusion. The captured images of these experiments contain five channels (Fig. 5a): nuclei stained with DAPI (blue), Titin-eGFP protein (green), Ttn-eGFP mRNA (red 570), mCherry-titin protein (red 610) and Ttn-mCherry mRNA (far red).

**Figure 5.**
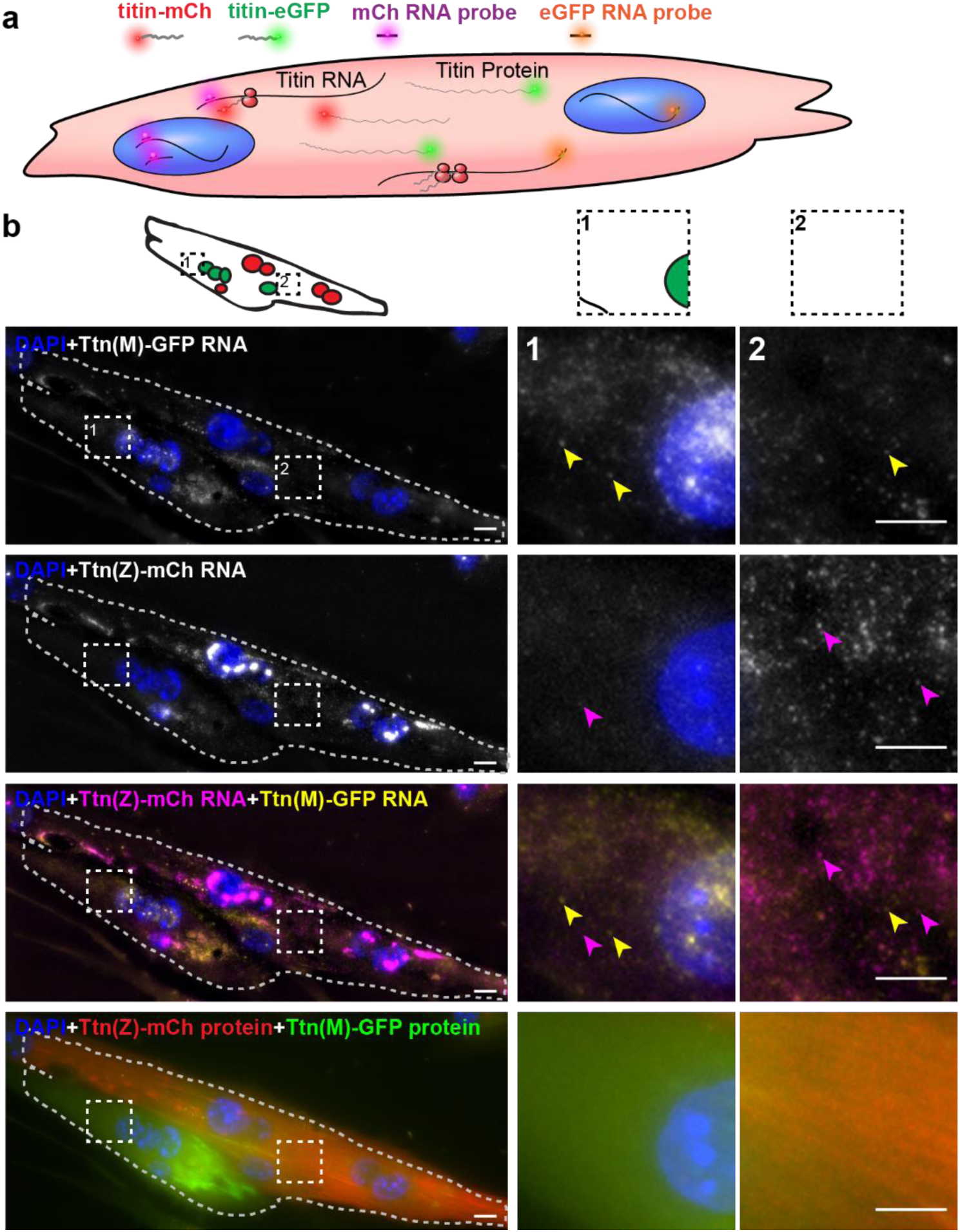
Distribution of titin mRNA in skeletal muscle cells undergoing cell fusion determined by smFISH detecting mCherry and GFP coding region. a) Model of titin mRNA and protein synthesis and localization. b) Representative image of a fusing myotube, where distribution of titin mRNA has just started. While we find both titin-mCh and titin eGFP mRNA in both the red and green compartments after cell fusion (magenta and yellow arrows), we do not see crossover of green and red titin fusion protein in the same area. Scale bar 10 µm.

Dots representing RNA signal were most intense in the nuclei and correspond to the transcription sites of titin (two main dots for two chromosomes in Fig. 5b). The nuclei contain only the mRNA from the cell they originated from, as confirmed by the strict separation of nuclear Ttn-mCherry or Ttn-eGFP mRNA. The myotube in the representative image of Fig. 5b had four nuclei with Ttn-eGFP (first row) and five nuclei with Ttn-mCherry mRNA (second row), summarized in the schematic overview above the image panel. The signal dots from Ttn-mCherry RNA appear much more intense than from Ttn-eGFP RNA signal dots and could relate to the insertion of mCherry at the 5’ end of the titin mRNA, which leads to an earlier transcription as compared to eGFP, inserted at the 3’ end. In the myotube in Fig. 5b, the titin proteins of different origin were not distributed completely over the whole syncytium (last row) indicating that fusion had just started. Therefore, there are still areas with mainly Ttn-eGFP protein (Fig. 5b magnification 1) or more mCherry-titin protein (magnification 2). In these areas we also found titin mRNA of both species, with mRNA from the distant nucleus underrepresented (e.g. titin-eGFP signal dots in the second magnification). In myotubes at a later stage after fusion with completely distributed titin protein (representative image in Fig. S4), titin mRNAs of both origins were present at the edge of the cell (magnification). These data indicate that it is not only titin protein which is distributed through the syncytium after cell fusion, but also titin mRNA.

### A theoretical approach to titin protein localization after cell fusion

We assume that red (green) titin is produced in the red (green) area and diffuses into the green (red) area while decaying according to the rate causing its half-life. The titin half-life in cultured skeletal muscle cells from day 12 chicken embryos is about 70 hours (Isaacs et al., 1989). In the adult mouse heart, tamoxifen induction of the conditional titin knockout leads to a maximum of ∼55% truncated titin after 80 days and ∼30% truncated titin after 5 days (Peng et al., 2007), suggesting a half-life of adult cardiac titin between 4 to 5 days (100 to 120h). Based on the embryonic chicken skeletal muscle and adult mouse heart data, we conservatively estimate the titin half-life at 3.5 days (τ=3.5d=3.024 10^5^ s). We estimated the titin diffusion coefficient D as 0.3 µm^2^s^-1^. The spatial decay length in a diffusion profile is (Dτ/0.693)^1/2^. The measured width of the titin gradient is d=50 µm (Fig. 4, 8h), which is not compatible with the τ and D values. If we accept the value for D, the value of τ required to explain this width is 0.693d^2^/D=1732.5µm^2^/0.3 µm^2^s^-1^ = 5775 s (<100 min), i.e. unrealistically short. If we accept the τ-value of 3.5 days, the diffusion coefficient to explain the gradient would be D=0.693d^2^/τ=5.7 10^-3^ µm^2^s^-1^, i.e. two orders of magnitude smaller than the value determined in cultured cells. Hence, another mechanism must act to restrict titin protein spread.

### Titin mobility and integration after *in vivo* regeneration and cell transplantation

The fusion of cultured myoblasts to multinucleated myocytes is a model for critical milestones in the development and regeneration of skeletal muscle. However, regeneration *in vivo* requires additional important steps and players, such as immune cells and the extracellular matrix. At the final stages of skeletal muscle formation *in vivo*, myotubes have formed muscle fibers, which are further differentiated and much larger than the myotubes that form *in vitro*. To evaluate if cell fusion provides additional benefits in animal experiments with cell transplantation (Darabi et al., 2012), we studied whether titin proteins from donor cells were distributed and integrated into the sarcomere lattice *in vivo*. Accordingly, we isolated donor myoblasts from Ttn(M)-eGFP mice and injected them into the tibialis anterior (TA) muscle of Ttn(Z)-mCherry mice one day after injury and induction of regeneration with cardiotoxin (CTX) (Experimental design Fig. 6a). Control groups received only CTX or only cell transplantation, respectively. After three weeks of regeneration, we dissected the treated and untreated contralateral TA muscles and cut them in half for longitudinal and transversal cryosections.

**Figure 6.**
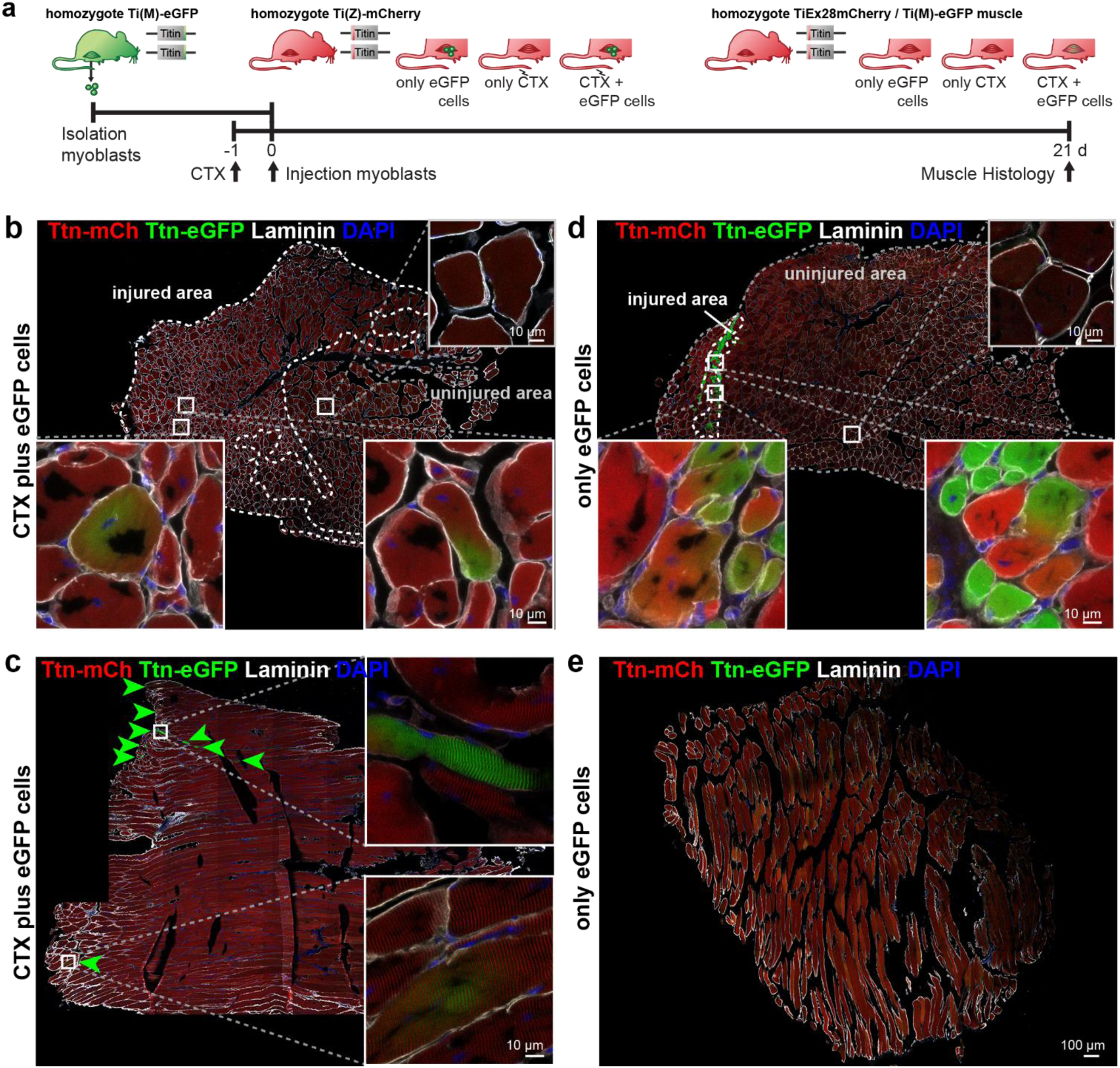
Titin distribution during regeneration. a) TA muscle of Ttn(Z)-mCherry mice was injured by injection of CTX followed by transplantation of Ttn(M)-eGFP myoblasts on the following day. Controls comprise CTX injury only and eGFP cell transplantation without injury. After 3 weeks of regeneration muscles (treated and untreated contralateral TA) were dissected and sections stained against laminin to visualize cell boundaries. Centralized nuclei in transversal sections (b, d) are a sign of regenerating cells within injured areas. These areas contain GFP positive fibers and extend throughout the muscle after CTX injury (b), but are limited to the injection site with cell injection only (d). The longitudinal sections (c, e) provide additional information about the proper integration of titin proteins from the transplanted cells into the regenerating myofibers. After injury, transplanted Ttn(M)-eGFP myoblasts fuse with the Ttn(Z)-mCherry cells of the injured host muscle and titin proteins from both cells contribute to the directionality of myofibers that is maintained along the muscle.

The injection of CTX caused muscle degeneration, followed by regeneration that was largely completed after three weeks, when individual regenerating cells were still present – as determined by their centralized nucleus (DAPI and laminin staining - Fig. 6b,d and S6a,c). In the control group with CTX only (Fig. S5a) and the mice with CTX and cell injection (Fig. 6b) the fibers contain mainly these centralized nuclei suggesting a successful completion of the degeneration-regeneration cycle. In the control group with only myoblast injection only very few myofibers contained centralized nuclei, located directly at the injection site (Fig. 6d). The untreated contralateral muscles had no fibers with central nuclei (Fig. S5c).

Successful transplantation of the Ttn(M)-eGFP myoblasts was detected in transversal sections with eGFP positive fibers in the injured area (Fig. 6b). In longitudinal sections we confirmed the proper integration of titin protein of the donor cells into the Ttn(Z)-mCherry muscle by the periodic staining of the myofilament in longitudinal sections (Fig. 6c). At several sites, muscle fibers were eGFP positive (green arrows, magnifications in Fig. S5e), as transplanted Ttn(M)-eGFP cells had differentiated together with the endogenous Ttn(Z)-mCherry satellite cells to mature muscle fibers. However, titin-eGFP signal was not evenly distributed over the complete fiber, but remained mainly proximal to the grafted nucleus.

Interestingly, in the control group with cell transplantation only (without prior injury) eGFP positive fibers were present at the injection site (Fig. 6d). Some of these fibers were also mCherry positive – primarily located along the injection canal. This finding is consistent with the insertion of the needle activating endogenous satellite cells, which subsequently fused with the transplanted eGFP myoblasts (Fig. 6d). In the control with CTX injury only (Fig. S6b) and in the contralateral muscles (Fig. S6d) eGFP positive fibers were absent.

In summary, cell transplantation can be used to deliver sarcomeric proteins to regenerating muscle. Without prior injury cells remained at the injection site (Fig. 6d,e), but in injured muscle donor cells distribute over a much larger area (Fig. 6b,c). Since eGFP-and mCherry-titin only intermix in a few, small fibers, diffusion of titin *in vivo* appears limited. Here, titin travels only in a limited area around the donor nucleus even after 3 weeks, so that a sarcomeric protein would be more confined to the fusion site versus the benefit of distributing the therapeutic protein over the whole syncytium.

## Discussion

Myofilament remodeling and adaptation is critical to balance efficient force generation and muscle mass. This includes how sarcomeres are formed, fortified, integrated into larger functional units and work in unison along the muscle fiber. Here, we take a visual approach towards understanding sarcomere and cell biology of skeletal muscle using a fluorescent mCherry-titin fusion protein (Z-disc label) expressed at physiological levels to complement the titin-GFP fusion protein (M-band label). Visualizing opposing sarcomere integration sites in double-heterozygous myocytes facilitates the analysis of sarcomere assembly and disassembly. We find increased mobility of Z-disk titin versus M-band titin in FRAP experiments. These data nicely complement our earlier work on cardiomyocytes (Rudolph et al., 2019). Most myotubes expressed at least two titin isoforms (biphasic fit of the fluorescent recovery curve), so that skeletal muscle cells appear more homogenous than cardiomyocytes with respect to titin isoform expression. Independent of the isoform makeup, protein exchange rates were largely similar between cardiac and skeletal muscle cells (Rudolph et al., 2019). Interestingly, the exchange is faster a the Z-disk than at the M-band, likely due to the integration of the newly synthesized protein with Z-disc titin mRNA available 1h earlier than M- band titin based on the speed of transcription (Jonkers and Lis, 2015). Alternatively, the contribution of short titins such as the Novex isoforms, which contain the Z-disk, but not the M-band sequences could help explain the difference.

We used the increased fluorescence of homozygous mCherry and eGFP knock-ins to study cell fusion and the outcomes of cell therapy as they allow the analysis of protein flux, compartmentalization, and the generation of functional units. Within hours after myotube fusion in cell culture, we found titin gradually distributed throughout the resulting syncytium. The spread of titin appeared to be facilitated in myotubes where mature sarcomeres had not yet formed. Nevertheless, even in myotubes that had already established a mature sarcomere structure, titin proteins of a newly fused cell were able to travel through almost half of the syncytium. Both protein and mRNA mobility contribute to the efficient distribution of titin after fusion.

Diffusion of proteins through the cytoplasm in myocytes versus non-muscle cells should be much more limited based on the high protein concentration in the cytoplasm and attachment to the dense cytoskeletal network within. The speed of diffusion is inversely correlated with the hydrodynamic radius of the protein (Arrio-Dupont et al., 1996) and packing titin in the myofilament structure limits protein diffusion even more. Microinjection of labelled dextran molecules into myotubes revealed the decrease of the diffusion coefficient with the molecular weight from 30 µm^2^/sec for a 9.5 kDa molecule to 2 µm^2^/sec for a 150 kDa molecule (Arrio-Dupont et al., 1996). In a similar experiment, globular proteins of different sizes were injected into isolated muscle fibers and diffusion coefficients differed depending on the fiber type likely due to differences in myofilament packing and not contraction (Papadopoulos et al., 2000). The distribution of titin along the myotube with about 1000 µm^2^/h (∼ 0.3 µm^2^/sec; Fig 4e) immediately after fusion, is relatively fast compared to the much smaller dextran molecules (Papadopoulos et al., 2000), suggesting a contribution of additional factors such as active transport (involving microtubules and the motor proteins kinesin or dynein) vs. passive diffusion.

The directed transport of mRNA to achieve proper subcellular localization is common in all types of cells and involves the interaction between Zip-code elements on the mRNA, multiple RNA-binding proteins and motor proteins. Thus, mRNA can be distributed 60 times faster than via passive diffusion and specific localization can be achieved. Transporting mRNA is more energy efficient than transporting protein, since many proteins can be translated from a single spatially organized mRNA (Buxbaum et al., 2015). Indeed myocytes use the scarce sarcomeric space to accommodate ribosomes even in adult muscle (Rudolph et al., 2019), suggesting that sarcomeric proteins are not transported actively in mature striated muscle cells, but rather produced on site from locally translated mRNA and limited distribution by diffusion.

In our cell culture model of myotube fusion, titin protein and mRNA from adjacent cells distribute throughout the sarcoplasm. Here, titin travels faster in cells without a mature sarcomere structure. In differentiated cells, sarcomeres are built from titins originating from both parental cells, resulting in an alternating striated pattern. Still, it has remained unclear if this also applies *in vivo*, where fusion events ultimately lead to large muscle fibers, which do not form *in vitro* (Almada and Wagers, 2016). To analyze how titin is distributed and integrated during regeneration and how healthy protein can be provided to diseased muscle *in vivo*, we used an injury model with injection of cardiotoxin into skeletal muscle (Garry et al., 2016) of the Ttn(Z)-mCherry mouse and transplanted Ttn(M)-eGFP myoblasts. As injury triggers the activation of the endogenous Ttn(Z)-mCherry satellite cells and their differentiation towards myocytes, myotubes and finally fibers, the transplanted Ttn(M)-eGFP cells differentiate as well and fuse with mCherry cells and fibers. Here, we found that fluorescent titin provides a strong label to not only quantitatively follow the repopulation of injured muscle with transplanted cells, but also evaluate the generation of a functional syncytium with continued directionality of myofibers. After 3 weeks of regeneration, eGFP positive fibers and their alternating fluorescent pattern confirmed the proper integration of donor titin. However, unlike in our tissue culture experiments, titin did not distribute throughout the fiber, but remained compartmentalized around the respective nucleus of origin. This might in part reflect the size difference between myotubes built *in vitro* from 2-10 cells and myofibers *in vivo* with up to hundreds of nuclei. *In vitro* generated fibers retained short mRNAs close to their nucleus, whereas long mRNAs like titin spread through the cell (Pinheiro et al., 2021), consistent with the titin mRNA localization in our smFISH experiments in myotubes. *In vivo*, single-nucleus RNA sequencing (sn-RNAseq) revealed also distinct nuclear subtypes and compartments (Kim et al., 2020), but did not allow statements of mRNA mobility. Our data would suggest that titin mRNA and the derived protein can cover distances of less than one millimeter, but will not travel from its nucleus of origin throughout the myofiber of several millimeters.

Ultimately, the difference between the fusion of cultured cells with homogenous distribution of titin versus compartmentalization of titin from donor cell and acceptor fiber in knock-in mice confirm the importance of *in vivo* studies towards understanding myocyte biology and extracting clinical relevance. Our mouse cell transplantation data suggest that in myopathies, compartmentalization of the therapeutic protein after fusion of a healthy cell with a diseased fiber might restrict the therapeutic effect (most prominent for the giant protein titin). To repopulate skeletal muscle with a relevant number of cells that deliver a therapeutic protein, it would therefore be beneficial to develop treatment protocols that target the early postnatal patient or consider in utero cell therapy approaches for a higher ratio of therapeutic to diseased cells and facilitated remodeling.

## Materials and Methods

### Generation of titin(Z)-mCherry knock-in mice

The mCherry cDNA was inserted into titin’s exon 28 (Z-disk) via a targeting vector (Fig. 1) using standard procedures (Radke et al., 2007). The animals were backcrossed on a 129/S6 background after successful integration.

### Genotyping

Genomic DNA was prepared from mouse ear biopsies with the HotSHOT method (Truett, 2000). The genotypes of the titin(Z)-mCherry (Primer: fwd CAGCATCATGGTAAAGGCCATCAA, rev CATTCAAATGTTGCCATGGTGTCC) and titin(M)-eGFP mice (Primer: AGAACAACAAGGAAGATTCCACA, AGATGAACTTCAGGGTCAGCTTG, TCTCAACCCACTGAGGCATA) were determined by PCR and visualized on agarose gels.

### Animal procedures

Mice were kept at the animal facility of the MDC in individually ventilated cages and a 12 h day and night cycle with free access to food and water. All experiments involving animals were performed according to institutional guidelines and had been approved by the local authorities (LAGeSo Berlin - Reg 0023/20). All surgeries were performed under isoflurane anesthesia, and every effort was made to minimize suffering. Strains are available upon request following institutional guidelines.

### Isolation and cultivation of primary myoblasts

For isolation of satellite cells from the titin(Z)-mCherry and titin(M)-eGFP lines, young mice (male and female) with an age of 3-4 weeks were used. Muscles from the hind limbs were collected and cut into small pieces. First digestion takes place by incubation in collagenase II (Sigma-Aldrich) for 30 min at 4°C followed by 20 min at 37°C. The second digestion step with collagenase/ dispase (Roche) is performed again first for 30 min at 4°C and then for 30 min at 37°C. The digestion is stopped and the tissue homogenate is filtered with 100 µm, 70 µm and 40 µm cell strainer. After centrifugation (1200 rpm, 10 min) the cells are resuspended in medium (DMEM/F12, 15% FBS, 50 µg/ml gentamicin, 1:1000 bFGF, 1:1000 LIF) and were pre-plated for 1-2 h to remove fibroblasts before they are seeded on matrigel (VWR) coated dishes. The cells were then cultivated complete medium (+ 1:50 B27) under 37°C and 5% CO2 and can be split or frozen.

The differentiation of the myoblast towards myotubes can be initiated by withdrawal of growth factors via changing to differentiation medium (DMEM, 5% horse serum, 1% Penicillin/streptavidin).

### Cardiotoxin injury and cell transplantation

For the analysis of titin integration and distribution during *in vivo* regeneration muscles of Ttn(Z)-mCherry mice were injured and myoblasts of Ttn(M)-eGFP mice were transplanted (n = 6 mice). Samples from three animals with insufficient myoblast integration were excluded from in depth analysis. The Ttn(Z)-mCherry mice were anaesthetized by isoflurane inhalation and the left TA muscle was injured by the injection of 40 µl of 10 µM cardiotoxin (CTX). Myoblasts of Ttn(M)-eGFP mice were isolated like described above and passaged two times before transplantation. 100 000 cells in 20 µl sterile PBS were injected into the left TA muscle one day after the injury. One mouse injected with CTX did not receive cell transplantation and served as a reference (CTX only control). Four mice received cell transplantation without the prior injury (cells only control). Adult male mice were block randomized, based on the litter, to the experimental or control groups. 21 days after the injury mice were euthanized and the treated and the untreated contralateral control TA muscles were dissected and fixed for histological analysis.

### smFISH

Cells were fixed with 2% PFA (sterile filtered) for 10 min at room temperature followed by permeabilization with 70% ethanol overnight at 4°C. The cells were then equilibrated in washing buffer (10% formamide and 2x saline sodium citrate (SSC) buffer) for 15 min at 37°C and the hybridization of the probes (100 nM in 10% formamide and 8% dextrane sulfate) with the target RNA was performed for 16 h at 37°C. DesignReady Stellaris® probe sets against mCherry (labelled with Quasar®-670, # VSMF-1031-5) and GFP (labelled with Quasar®-570, # VSMF-1014-5) from Biosearch Technologies were used.

After washing the cells for 30 min at 37°C, they were stained with DAPI (1:2000 in washing buffer) for 10 min at 37°C and washed with 2x SSC buffer. The imaging was performed directly on the next day to prevent degradation of the RNA.

The samples were imaged with a widefield microscope (Nikon Eclipse Ti) with narrow Bandpass filter and a 63x objective. They were excited with the Prior Lumen 200 system and the following filters were used: DAPI (Ex: 387/11, Em: 447/60, beam splitter: HC BS 409), GFP (Ex: 470/40, Em: 525/50, BS: T 495 LPXR), Quasar®-570 (Ex: 534/20, Em: 572/28, BS: HC BS 552) and CalFluor®-610 (Ex: 580/25, Em: 625/30, BS: T 600 LPXR), Quasar®-670 (Ex: 640/30, Em: 690/50, BS: T 660 LPXR). 21 z-stack images with 0.3 µm steps were taken with the Andor DU888 camera. Images were processed with the Fiji (Fiji is just ImageJ) software. Background was reduced for the mRNA channels by subtraction with a Median filtered (50 px) copy of the image and z-stacks were projected with maximal intensity.

### Immunofluorescence staining

TA and EDL muscles were dissected, fixed with 4% PFA, dehydrated in 30% sucrose and frozen in Tissue-Tek® O.C.T.^TM^. Cryosections of these tissues were performed with a thickness of 10 µm. The sections were permeabilized and blocked with blocking solution (10% goat serum, 0.3% Triton X 100 and 0.2% BSA in PBS) for 2 h. Cells were fixed with 4% PFA at room temperature for 10 min and washed with PBS followed by permeabilization and blocking as above. The incubation with the primary antibody (diluted in PBS) was performed at 4°C overnight (α-actinin (A7811, Sigma, RRID:AB_476766) 1:100, Laminin (L9393, Sigma, RRID:AB_477163) 1:100, M-Cadherin (sc-81471, SantaCruz, RRID:AB_2077111) 1:50). After washing five times with PBS, cells were incubated with a fluorescent secondary antibody (diluted 1:1000 in PBS) for 2 h at room temperature. Stained sections and cells were mounted with ProLong Gold mounting medium.

Confocal images were acquired with a laser-scanning microscope (LSM700 and LSM710, Carl Zeiss) with a Plan-Apochromat 63x/1.4oil Ph3 objective or a Plan-Apochromat 20x/0.8 M27 objective for overview images. Qualitative images were replicated at least three times and representative images were shown. Line profiles were created out of the raw, unmodified images using the Fiji software and fluorescence intensity was normalized.

### Live Imaging

Live imaging experiments were carried out on the DeltaVision Elite microscope (GE healthcare) or the CSU-W1 SpinningDisk (Nikon) microscope. For the DeltaVision microscope, the 60x oil objective (NA 1.42) was used with the FITC filter set for imaging eGFP and the A594 filter set for mCherry imaging. The 40x objective (NA 1.15) was used for the SpinningDisk microscope and a GFP and a mCherry filter set. The incubator of the microscopes was adjusted and equilibrated to 37°C and 5% CO_2_ prior imaging and a humidifier was used. Cells were kept in FluoroBrite medium (plus identical supplement as during cultivation) during imaging. To avoid photo-toxicity, the laser powers were adjusted as low as possible. Usually several cells (about ten) were selected in a point list and imaged every 30-60 min for 12-16 h at five z-stacks. To avoid shifting of the focus during the hours of imaging, the UltimateFocus option of the DeltaVision and the Perfect focus system of the SpinningDisk microscopes were used.

The imaging of fusing myotubes areas with red and green cells in close proximity were selected. Since it was expected that only a part of these cells fuse during the selected time span, many areas (about 20 to 30) were selected in each experiment. The progression of fusion was measured by selecting ROIs based on the fluorescence intensity threshold. As a first step, the fluorescence intensity of a negative and a bright positive neighboring cell was measured and set as 0 and 100%, respectively. The fluorescence intensity value representing 20% was selected as first threshold (weak signal) and the 50% value as second threshold (strong signal). These thresholds were used to define regions with no, weak or strong signal for red and green. In the fusion process, the overlap of red and green signals of different intensities were used to assign 5 different types of ROIs:

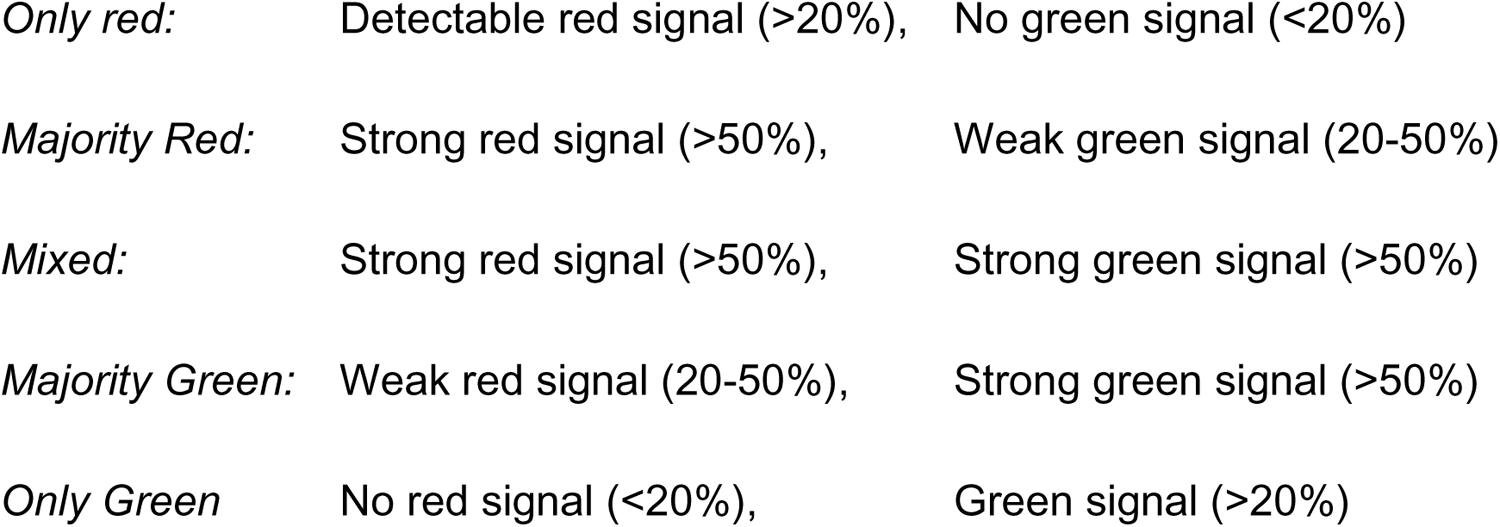

### Fluorescence recovery after photobleaching

FRAP experiments were performed on the DeltaVision Elite microscope with the 60x oil objective (NA 1.42). Both fluorophores of the double-heterozygous Ttn(Z)-mCherry/Ttn(M)-eGFP myotubes were photobleached with a 488 nm laser at 25% intensity for 0.1 s. A rectangular region of interest (ROI) covering two sarcomeres is bleached and the fluorescence recovery was followed over 14 h with imaging every 5 min for the first 30 min, then every 30 min for another 1.5 h and every hour for the last 12 h. Three individual experiments with three cells each were performed. Fluorescence intensity was measured at the respective integration sites and between it. The signal intensities were normalized to the intensities before bleaching and to the intensities of the whole cell like it is described by (Al Tanoury et al., 2010).

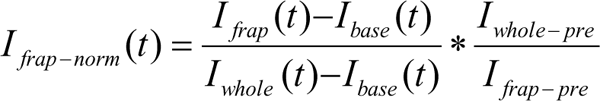

These normalized data were then used to fit a one-phase association curve to it with GraphPad Prism.

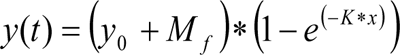

This curve was then used to calculate the exchange half-life, which is the time point when 50% of the maximal signal has recovered.

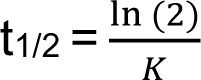

Most of the myotubes exhibited a recovery kinetics that could be fitted better with a two-phase association curve:

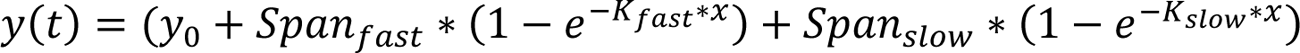

This biphasic recovery is divided in fast and a slow phase. With this formula, GraphPad Prism also calculates the percentage of the fast phase.

Independent of the type of recovery, the mobile fraction can be calculated by the fluorescence intensity at the end (when the plateau is reached) relative to the intensity at the beginning (da Silva Lopes et al., 2011).

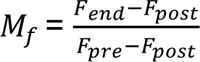

### Statistics

Statistical analysis was done with the GraphPad Prism Software (Version 5). Differences between two data sets are analyzed by t-test and differences between three or more data sets with one-way ANOVA. Data affected by two factors are analyzed by two-way ANOVA and Bonferroni posttest. Significances are indicated with *, P < 0.05; **, P < 0.01; ***, P < 0.001. Number of biological replicates are indicated in the respective figure legends.

## Acknowledgement

This work was funded by the European Research Council (ERCAdv to M.G.) and the German Research Foundation (DFG to M.G.) and by DZHK (German Centre for Cardiovascular Research)-project MD3-Nanopathology (to S.E.L.). We thank Anje Sporbert, Anca Margineanu and the Microscope Core Facility from the MDC for support with the confocal microscopes, and Janine Fröhlich for expert technical assistance.

## Author contributions

Judith Hüttemeister and Franziska Rudolph planned and performed experiments and analyzed the data supported by Claudia Fink. Michael Radke generated the mouse model and conducted animal experiments. Martin Falcke supported imaging data interpretation. Eva Wagner and Stephan Lehnart provided access to technology. Dhana Friedrich and Stephan Preibisch provided access to technology and supported the smFISH experiments. Michael Gotthardt and Judith Hüttemeister wrote the manuscript with input from all authors.

## Material & Correspondence

Correspondence and material requests should be addressed to M.G.

## Supplement

### Supplemental figures

**Figure S1.**
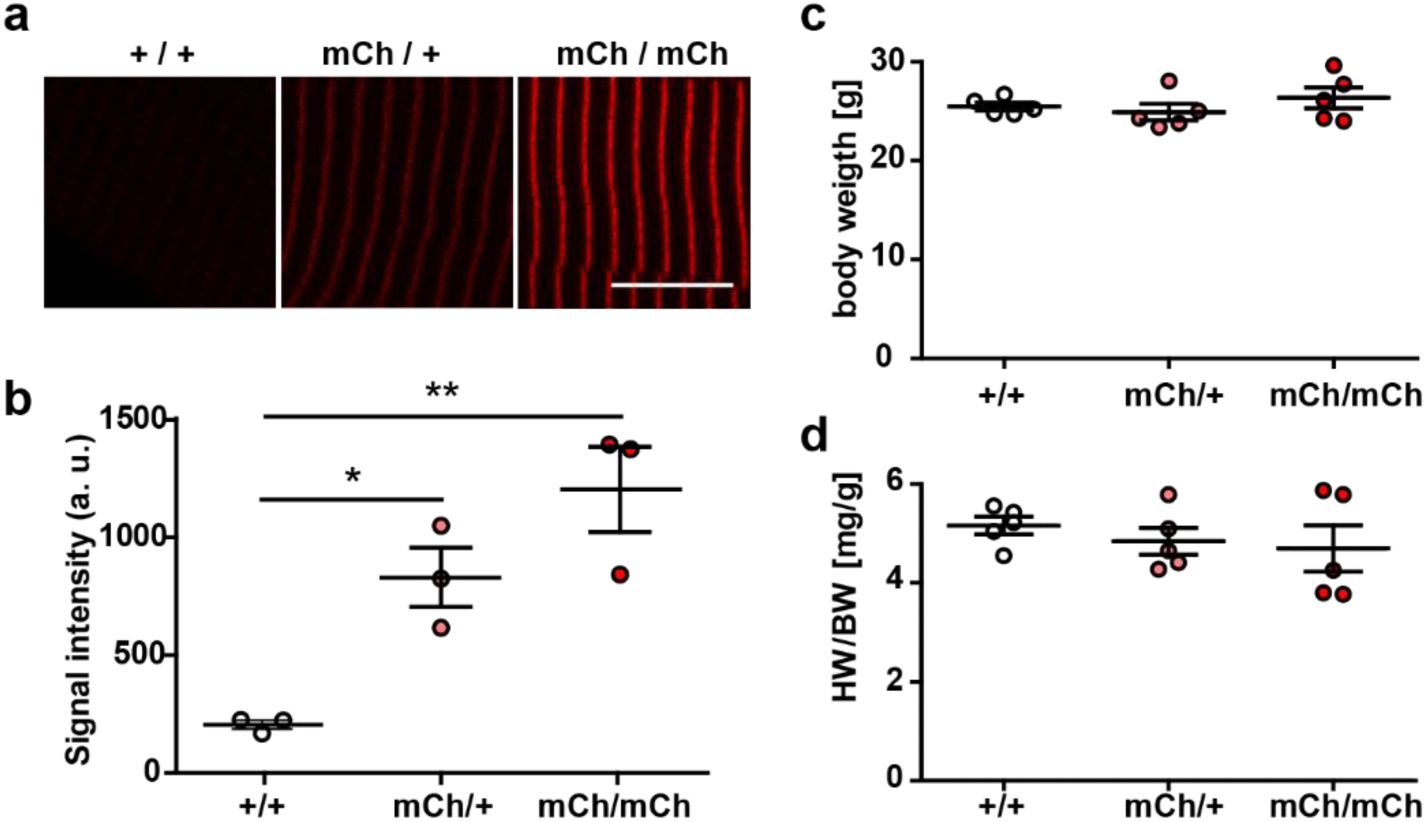
Knock-in of mCherry at titin’s Z-disk. a) Representative images of extensor digitorum longus (EDL) sections of homozygous, heterozygous and WT Ttn(Z)-mCherry mice with significantly different fluorescence intensities (b); one-way ANOVA (n=3 with 5 sarcomeres per replicate). Body weight (c) and heart to body weight ratio (HW/BW, d) were not significantly different between genotypes. Scale bar 10 µm.

**Figure S2.**
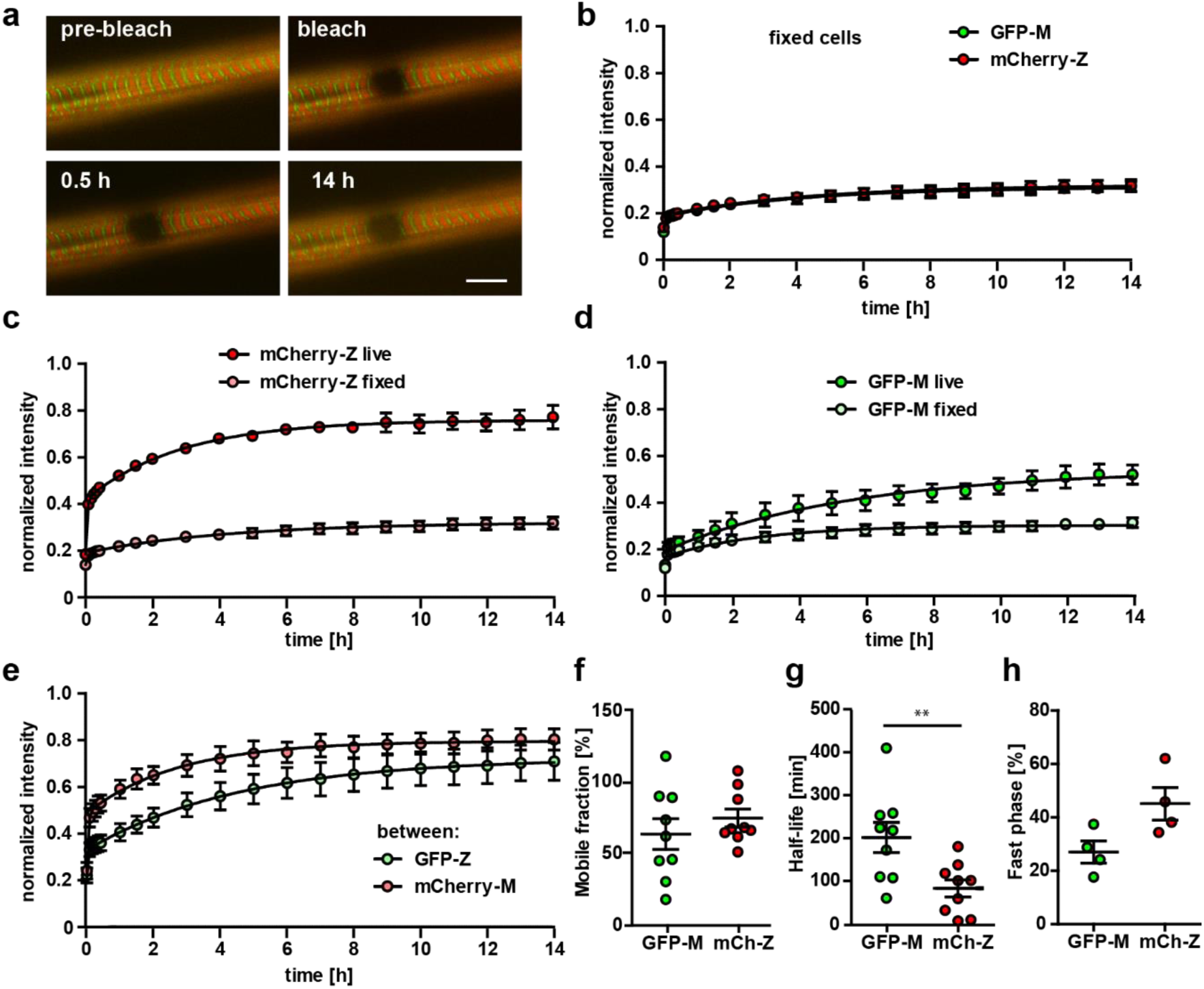
Titin kinetics in double-heterozygous myotubes. a) Representative images of FRAP in fixed cells reveal no recovery of titin signal over 14 h at Z-disk and M-band. Scale bar 10 µm. b) No difference in the reactivation of the fluorophore for GFP and mCherry, but highly significant difference between the recovery in live and fixed cells (c, d). The difference in recovery between mCherry labelled Z-disk titin and GFP-labeled M-Band titin is not as pronounced outside their respective integration sites (e). For the titin kinetics outside the integration site measured by mobile fraction (f), exchange half-life (g) and fast phase (h), only half life is significantly reduced between Z-disk and M-band label.

**Figure S3.**
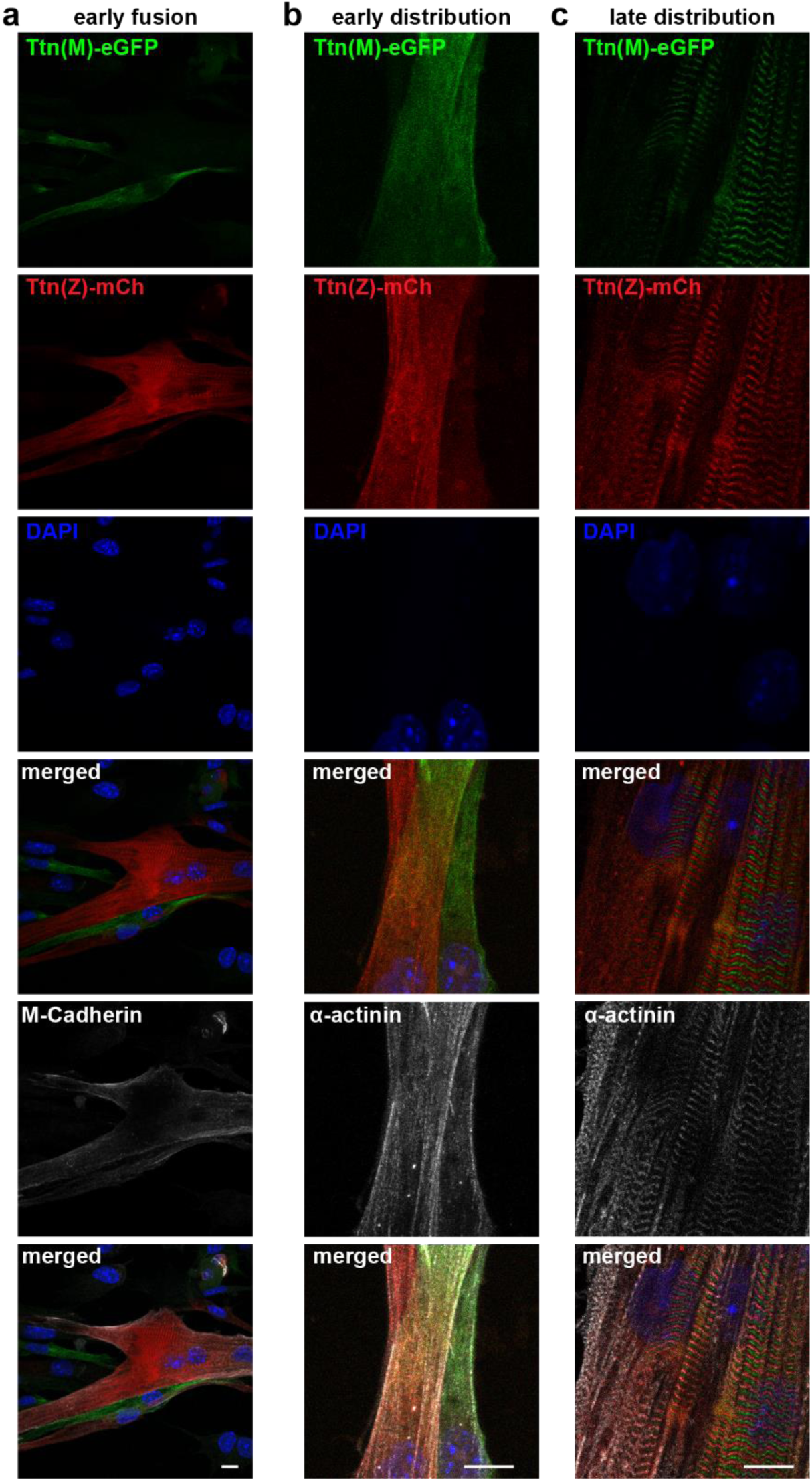
Titin distribution after cell fusion. Satellite cells isolated from Ttn(Z)-mCherry and Ttn(M)-eGFP animals were co-cultured, fixed at different fusion states (a: early fusion, b: early distribution, c: late distribution) and co-stained against M-cadherin or α-actinin. Titin is distributed throughout the fused cells, at early and late stage. Scale bar 10 µm.

**Figure S4.**
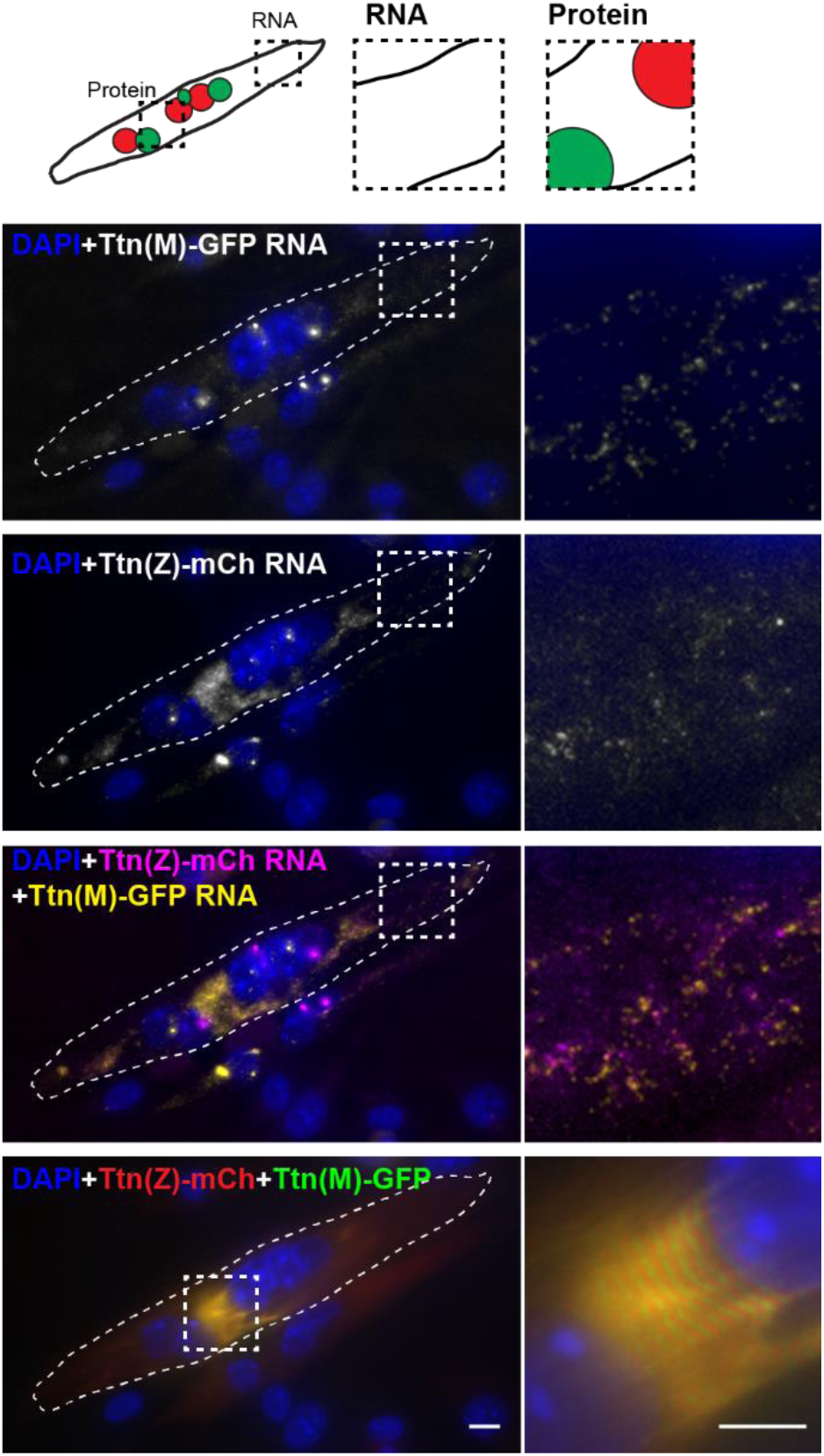
Titin mRNA localization after cell fusion. Satellite cells were isolated from homozygous Ttn(Z)-mCherry and Ttn(M)-eGFP mice, co-cultured, differentiated to myotubes and fixed after fusion. The cartoon indicates the position of nuclei expressing mCherry vs. GFP titin fusion proteins. Representative image of a fused myotube with intermixed protein and RNA and mature sarcomere structure. Scale bar 10 µm.

**Figure S5.**
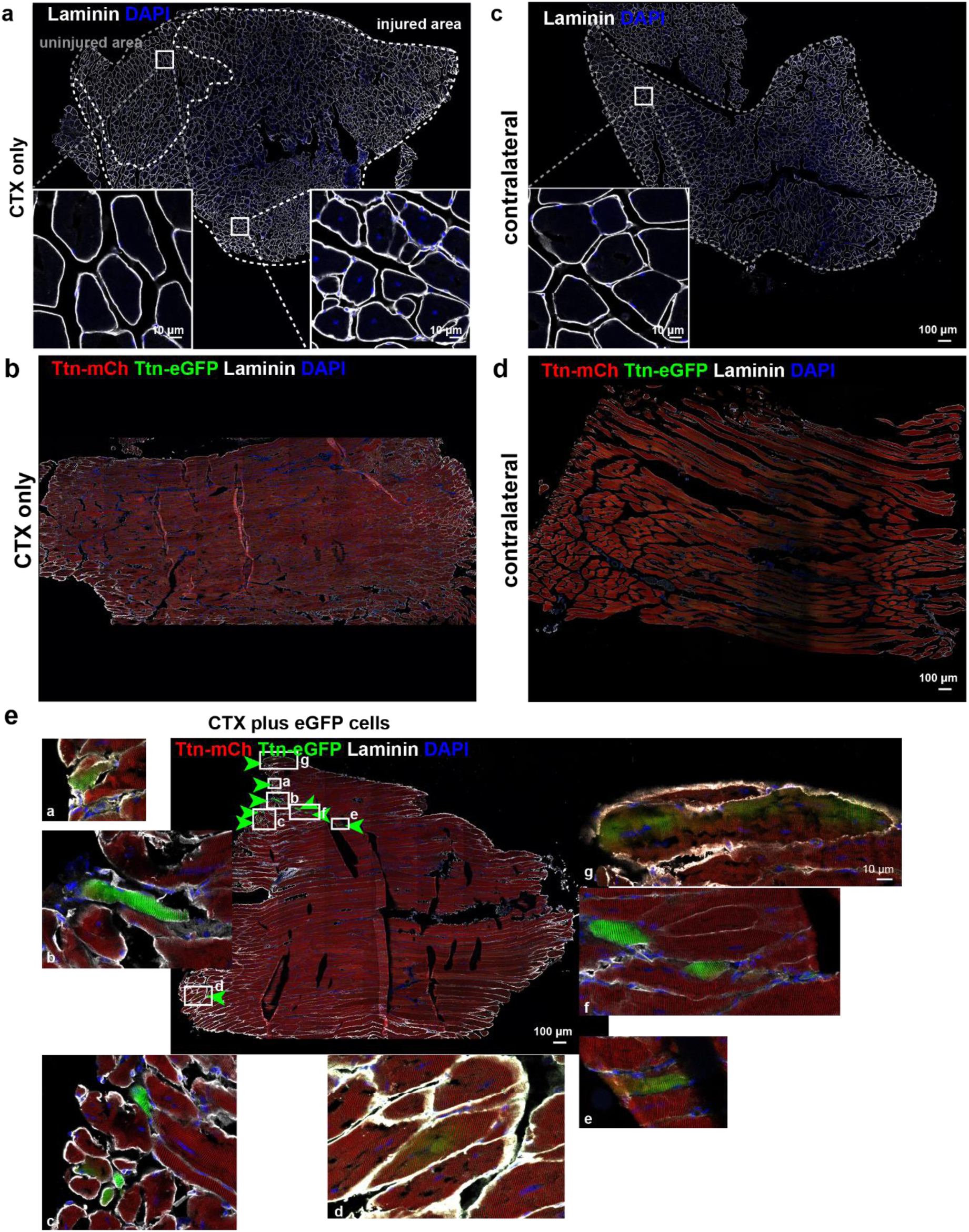
Titin mobility and integration upon *in vivo* regeneration and cell transplantation. a) Transversal sections of tibialis anterior (TA) muscles of the control group (CTX only) and the non-injected contralateral muscle (b). c) Longitudinal sections of TA muscles of the CTX only group and the contralateral muscle (d). e) Magnifications of areas with eGFP positive fibers from the longitudinal section of the CTX injury plus cell transplantation group (white letters a to g for seven regions of interest). Scale bar 100 µm.

